# Single-cell transcriptomics reveals the identity and regulators of human mast cell progenitors

**DOI:** 10.1101/2021.10.01.462521

**Authors:** Chenyan Wu, Daryl Boey, Oscar Bril, Jennine Grootens, M. S. Vijayabaskar, Chiara Sorini, Maria Ekoff, Nicola K. Wilson, Johanna S Ungerstedt, Gunnar Nilsson, Joakim S. Dahlin

**Author notes:** Corresponding author Joakim Dahlin; Address: NKS BioClinicum J7:30, Visionsgatan 4, 171 76 Solna, Sweden. These authors contributed equally (Chenyan Wu, Daryl Boey).

## Abstract

Mast cell accumulation is a hallmark of a number of diseases including allergic asthma and systemic mastocytosis. IgE-mediated crosslinking of the FcεRI receptors causes mast cell activation and contributes to disease pathogenesis. The mast cell lineage is one of the least studied among the hematopoietic cell lineages and there are still controversies about the identity of the mast cell progenitor, i.e., whether FcεRI expression appears during the hematopoietic progenitor stage or in maturing mast cells. Here, we used single-cell transcriptomics to reveal a temporal association between the appearance of FcεRI and the mast cell gene signature in CD34^+^ hematopoietic progenitors. In agreement with these data, the FcεRI^+^ hematopoietic progenitors formed morphologically, phenotypically and functionally mature mast cells in long-term culture assays. Single-cell transcriptomics analysis further revealed the expression patterns of prospective cytokine receptors regulating mast cell progenitor development. Culture assays showed that IL-3 and IL-5 promoted disparate effects on progenitor cell proliferation and survival, respectively, whereas IL-33 caused robust FcεRI downregulation. Taken together, we have demonstrated that FcεRI appears during the hematopoietic progenitor stage of mast cell differentiation and that external stimuli may regulate the FcεRI expression. Thus, the results resolve the controversy regarding the appearance of FcεRI during mast cell development.

**One-sentence summary:** Single-cell analysis of human hematopoiesis uncovers the stage at which FcεRI appears during mast cell differentiation and reveals disparate effects of IL-3, IL-5 and IL-33 on mast cell progenitor proliferation, survival, and suppression of FcεRI expression.

## Introduction

Mast cells contribute to the pathogenesis of various diseases including allergies, asthma, and mastocytosis (*1*). One of the mast cells’ main responses follows IgE-allergen immune complex crosslinking of the high-affinity IgE receptors, FcεRI, on the cell surface. This event leads to mast cell activation and degranulation, ultimately causing an allergic response. The signaling that follows FcεRI crosslinking has been studied in detail. However, the regulation of the FcεRI expression during human mast cell development is unclear and has remained a disputed subject.

The consensus is that mast cell progenitors exhibit low, if any, FcεRI expression (*2*). This view favors a mechanism in which FcεRI expression is upregulated during terminal mast cell maturation (*3*). Evidence includes in vitro-based cell culture experiments, which show that FcεRI expression appears following the decrease of the progenitor marker CD34 (*4, 5*). Existence of FcεRI^−^ and FcεRI^+^ primary mast cells in asthmatic lungs also supports the view that FcεRI is a late event in mast cell maturation (*6*). By contrast, some studies present an opposing view and suggest that FcεRI expression may appear already at the progenitor stage. For example, short-term culture of FcεRI^+^ hematopoietic progenitors in multi-cytokine conditions produces cells with mast cell-like characteristics (*7*). However, available studies have not cultured FcεRI^+^ hematopoietic progenitors in mast cell-promoting conditions for an extended period of time (*7, 8*), which is typical for in vitro-derived mast cells (*9*). Thus, it has been easy to disregard findings that potentially indicate mast cell-forming potential of FcεRI^+^ hematopoietic progenitors.

The present investigation aimed to study the appearance of FcεRI during mast cell development in human hematopoiesis. We performed single-cell RNA sequencing with an extended cell hashing-based approach to generate a transcriptional landscape of hematopoietic progenitor differentiation in peripheral blood. A mast cell gene signature was associated with upregulation of FcεRI at the gene and protein level. In agreement with this, sorted and long-term cultured FcεRI^+^ progenitors formed cells that phenotypically, morphologically, and functionally exhibited features of mast cells. Notably, single-cell transcriptomics analysis and cell culture experiments identified IL-33 as a modulator of FcεRI expression on mast cell progenitors. Taken together, FcεRI^+^ hematopoietic progenitors exhibit mast cell-forming potential but microenvironmental conditions are able to downregulate FcεRI expression.

## Results

### Single-cell transcriptomics reveals blood cells expressing mast cell genes and FcεRI subunit genes

The developmental stage at which mast cells upregulate the FcεRI expression has yet to be determined. To investigate whether FcεRI appears early in mast cell differentiation, we performed single-cell RNA sequencing analysis of cells isolated from healthy peripheral blood, in which mature mast cells are absent. Sorting Lin^−^ c-Kit^+^ cells allowed us to profile a spectrum of differentiating hematopoietic progenitor cells using droplet-based single-cell RNA sequencing. We visualized the 6874 single-cell transcriptomes in two dimensions using UMAP (Fig 1A). Plotting established marker genes for various lineages enabled annotation of the main branches and populations of the hematopoietic landscape (Fig 1A, Fig S1) (*10-13*). We also noted a distinct cluster of cells with a mast cell gene signature (*TPSAB1, TPSB2*, and *HDC*), hereon referred to as cluster A (Fig 1A-B). Notably, cells in cluster A expressed the genes *FCER1A, MS4A2*, and *FCER1G* – coding for the three protein subunits that form FcεRI.

**Fig 1.**
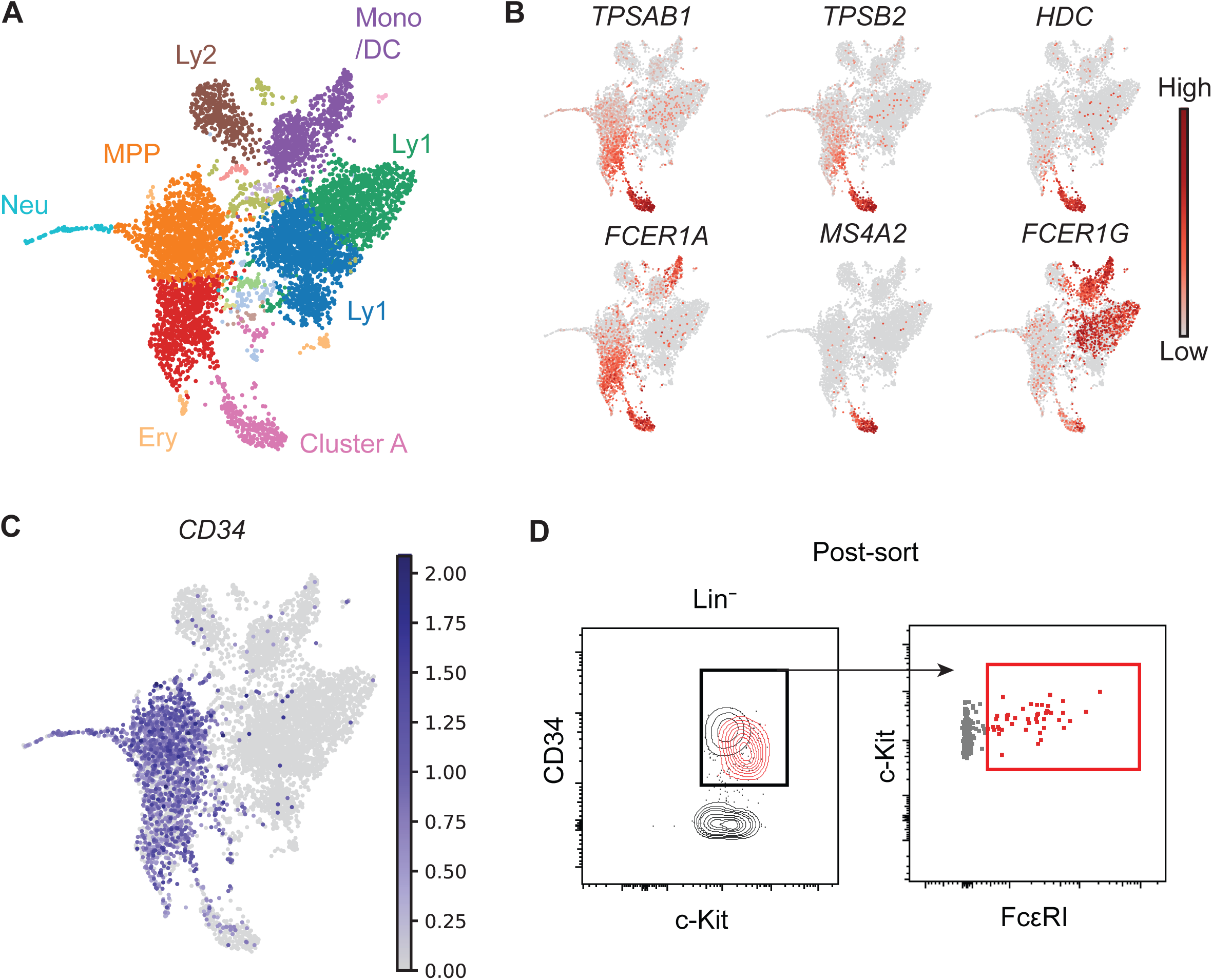
Expression of Fc Ε RI subunit genes in hematopoietic progenitors is associated with a mast cell gene signature. **A**) UMAP visualization of the single-cell transcriptomes of 6874 Lin^−^ c-Kit^+^ peripheral blood cells. The colors indicate Leiden clusters. Expression patterns of lineage genes used to annotate the landscape are shown in Fig S1. **B**) Gene expression levels of mast cell genes and Fc Ε RI subunit genes. **C**) Gene expression of *CD34*. **D**) Post-sort flow cytometry analysis of Lin^−^ c-Kit^+^ cells.

We next devised a plan to generate a second single-cell RNA sequencing dataset that captures the spectrum of hematopoietic progenitors and cells with the mast cell-like phenotype at high resolution. To determine the isolation strategy of this prospective dataset, we first plotted *CD34* expression on the Lin^−^ c-Kit^+^ landscape, visualizing the hematopoietic progenitor cells (Fig 1C). Cluster A cells exhibited residual *CD34* expression, which indicated that these cells also had an immature phenotype. Flow cytometric analysis verified the presence of a CD34^+^ FcεRI^+^ subpopulation among sorted Lin^−^ c-Kit^+^ cells (Fig 1D). These cells exhibited lower CD34 protein expression than the bulk of the CD34^+^ cell population, which agrees with the low *CD34* gene expression levels in cluster A. The results suggested that the generation of a Lin^−^ CD34^+^ c-Kit^+^ dataset would capture a spectrum of hematopoietic progenitor cells and include Lin^−^ CD34^+^ c-Kit^+^ FcεRI^+^ mast cell progenitor-like cells, hereon referred to as population A.

### Single-cell transcriptomics reveals an association between FcεRI expression and a mast cell gene signature in hematopoietic progenitors

Population A is a subset of the larger Lin^−^ CD34^+^ c-Kit^+^ cell fraction (Fig 2A). However, retrospective identification of population A cells within a Lin^−^ CD34^+^ c-Kit^+^ single-cell transcriptomics dataset is challenging, as population A is defined based on surface protein phenotype. We took advantage of a cell hashing-based method for single-cell RNA-sequencing to generate a single comprehensive dataset of hematopoietic progenitors with additional enriched (spiked-in) barcoded population A cells (*14*). Briefly, dividing the sample of cells before FACS isolation allowed labelling of a fraction of the cells with phycoerythrin (PE)-oligonucleotide barcodes, which can be detected by the single-cell RNA sequencing protocol (Fig 2B). We subsequently sorted PE-oligonucleotide-labeled population A cells, i.e. Lin^−^ CD34^+^ c-Kit^+^ FcεRI^+^ cells, and pooled these cells with unlabeled hematopoietic progenitors (Fig S2). Flow cytometry analysis of the cell pool revealed a bimodal distribution of the PE-oligonucleotide, which was suggestive of successful barcoding, and showed that 29 % of the cells represented spiked-in population A cells (Fig 2C). Single-cell RNA sequencing with feature barcoding technology allowed simultaneous measurement of gene expression and oligonucleotide levels of the pooled cells. Visualization of the entire single-cell transcriptomics dataset using UMAP revealed the erythrocyte and neutrophil entry points (Fig 2D and Fig S3). A branch of cells expressing mast cell signature genes and FcεRI subunit genes was also present (Fig 2E). To verify whether these cells constituted the population A cells, we first plotted a histogram of the hashtag oligonucleotide levels of each single cell (Fig 2F). The hashtag oligonucleotide levels showed a bimodal distribution pattern, representing unlabeled hematopoietic progenitor cells and oligonucleotide-labeled population A cells. Notably, the ratio between the spiked-in oligonucleotide-labeled population A cells and the hematopoietic progenitors was similar between the flow cytometry and single-cell transcriptomics analysis (Fig 2C,F). Visualization of the oligonucleotide-labeled cell fraction in the hematopoietic landscape established the location of population A, which was divided into two main clusters (Fig 2D,G).

**Fig 2.**
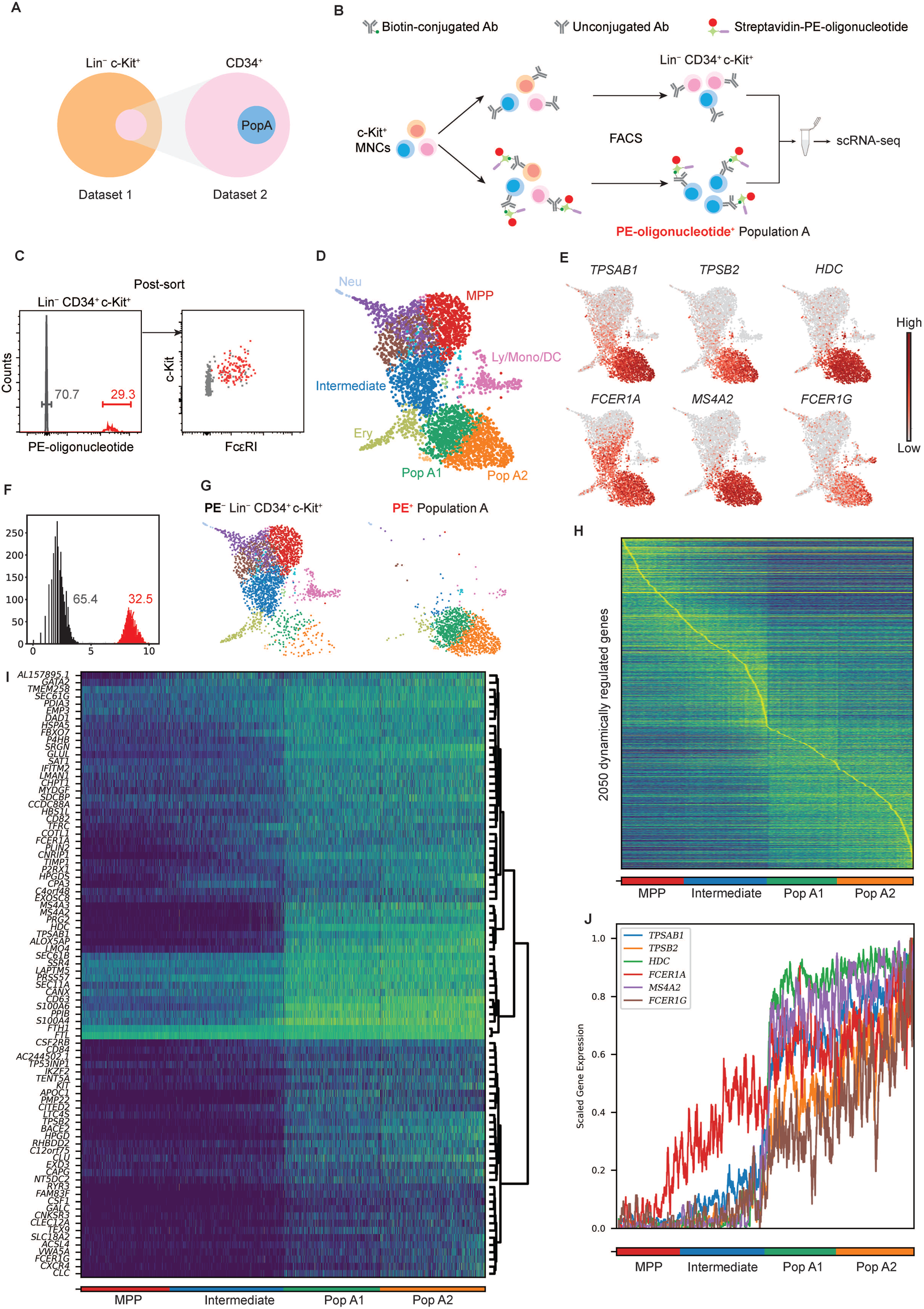
Single-cell transcriptomics reveals a temporal association between Fc Ε RI and the mast cell gene signature. **A**) Representation of the cells in the first dataset relative to the cells in the prospective second dataset. **B**) Experimental approach to hashtag a cell subset with a PE-oligonucleotide barcode. **C**) Post-sort flow cytometry analysis of pooled cells, constituting Lin^−^ CD34^+^ c-Kit^+^ cells without and Lin^−^ CD34^+^ c-Kit^+^ Fc Ε RI^+^ cells with the PE-oligonucleotide hashtag. **D**) UMAP visualization of the entire Lin^−^ CD34^+^ c-Kit^+^ dataset (4063 cells). The colors indicate Leiden clusters. See panels E, G and Fig S3 for landscape annotation. Population A consisted of two main clusters, Pop A1 and Pop A2. **E**) Gene expression levels of mast cell genes and Fc Ε RI subunit genes. **F**) Histogram showing the distribution of the PE-oligonucleotide in the single-cell transcriptomics data. Grey bins represent ambiguous events. **G**) Plots showing the PE^−^ Lin^−^ CD34^+^ c-Kit^+^ cells and PE^+^ Lin^−^ CD34^+^ c-Kit^+^ Fc Ε RI^+^ cells (population A) **H**) Visualization of highly dynamic genes along the mast cell differentiation trajectory. Cells are ordered according to cluster and pseudotime. **I**) Visualization of dynamic genes that positively correlate with rank ordering, also referred to as Pop A signature genes. **J**) Plot showing the temporal upregulation of mast cell genes and Fc ΕRI subunit genes. Ery, erythroid; MPP, Multipotent progenitor; Ly/Mono/DC, lymphocyte/monocyte/dendritic cell; Neu, neutrophil; Pop A, population A.

To study the temporal association between population A signature genes and FcεRI subunit genes, we reconstructed the differentiation trajectory from multipotent progenitor to population A cells. Dynamically regulated genes (*15*) were identified by leveraging cell clustering and diffusion pseudotime inferences (Fig 2H). Correlation analysis between gene expression levels and differentiation stage gave rise to a gene signature associated with population A (Fig 2I). The signature genes of population A included genes involved in the production of proteases (*CPA3, TPSAB1, TPSB2*), granule-associated mediators (*HDC, SRGN*), FcεRI (*FCER1A, MS4A2, FCER1G*), cell differentiation and survival (*KIT*), cell activation (*CD63*), and prostaglandin-associated enzymes (*HPGDS, HPGD*). Of note, 9 out of 14 well-defined mast cell genes were represented within population A signature genes (Table S1) (*16*).

We next focused on the temporal upregulation of hallmark mast cell genes and their relation to FcεRI subunit genes during the differentiation from multipotent progenitors to population A cells. Plotting the gene expression along the trajectory revealed that upregulation of the mast cell genes *TPSAB1, TPSB2*, and *HDC* sharply increased upon differentiation into early population A cells (Fig 2J). *FCER1A*, which codes for the α subunit of FcεRI, was expressed in progenitors preceding the population A stage. However, mast cells express all three FcεRI subunits – the α, β and γ subunits. Notably, the γ subunit is required for surface expression of the FcεRI protein (*17, 18*), and the corresponding gene, *FCER1G*, is upregulated together with the mast cell genes. Taken together, the single-cell transcriptomics analysis shows an association between FcεRI and the appearance of a mast cell gene signature.

### Hematopoietic progenitors expressing FcεRI rapidly form mast cells in vitro

The association between the mast cell signature and the FcεRI subunit genes in CD34^+^ progenitors prompted us to investigate whether population A exhibited mast cell-forming potential. We FACS-sorted and cultured population A progenitors, using FcεRI^−^ progenitors as a reference. The isolated progenitors were long-term cultured in mast cell-promoting conditions (Fig 3A), similar to previous studies (*19, 20*).

**Fig 3.**
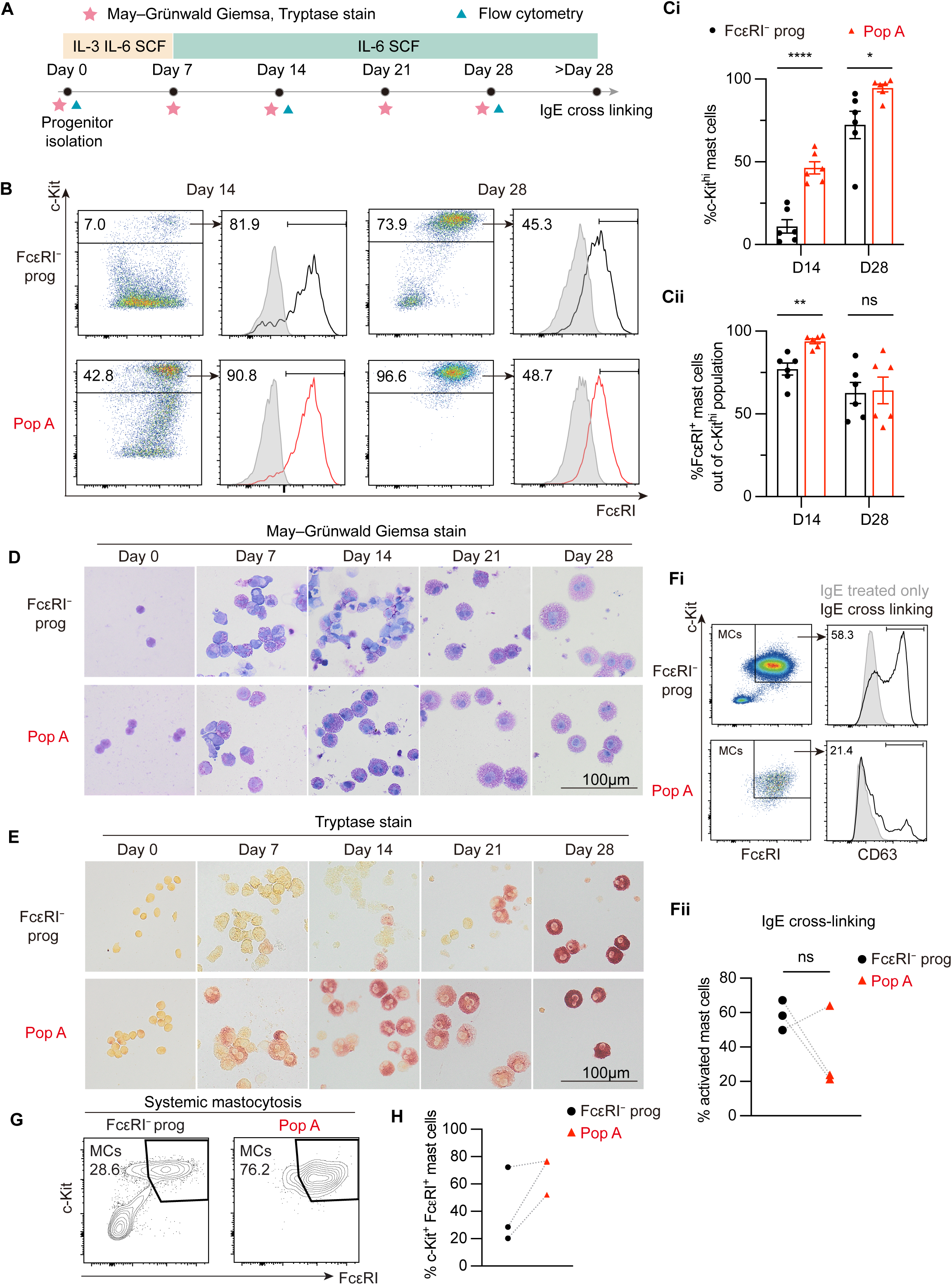
Fc ΕRI^+^ hematopoietic progenitors exhibit mast cell-forming potential in health and disease. A) Experimental setup to analyze mast cell differentiation. 10 ng/ml IL-3, 50 ng/ml IL-6, 100 ng/ml stem cell factor (SCF). B-C) Flow cytometry analysis of cultured progenitors (n=6). Filled histograms show isotype control staining. D) May-Grünwald Giemsa staining. E) Tryptase staining (day 0, one experiment; days 7-28 from an individual experiment, culture experiment performed 6 times). Fi) Functional analysis of mast cells derived from hematopoietic progenitors. The CD63 gate that defines mast cell activation was set to ensure that the IgE only group showed less than 1 % positive events. Fii) Quantification of activated mast cells in panel Fi (n=3). Dotted lines link samples from the same donor. G) Flow cytometry analysis of cells derived from patients with systemic mastocytosis. H) Quantification of the results in panel G (n=3). Dotted lines link samples from the same donor. Two-tailed unpaired t-tests, \**P*<0.05, \*\**P*<0.01, \*\*\*\**P*<0.0001.

A combination of flow cytometry, cytochemical staining, and enzymatic staining to assess tryptase activity was used to establish the phenotype following long-term culture (Fig 3A). The progenitors, irrespective of initial FcεRI expression, demonstrated the potential to form c-Kit^hi^ mast cells (Fig 3B-C). However, the flow cytometry analysis revealed that population A progenitors exhibit a more rapid progression into the c-Kit^hi^ mast cells phenotype compared with FcεRI^−^ progenitors. Similarly, morphological analysis revealed that population A progenitors quickly formed granulated tryptase-expressing mast cells (Fig 3D-E). Crosslinking the FcεRI receptors on the resulting mast cells caused CD63 upregulation on the cell surface (Fig 3F).

It has previously been proposed that mast cells from mastocytosis patients originate from CD34^+^ FcεRI^−^ progenitors (*5*). We isolated population A cells from three patients with systemic mastocytosis to investigate whether FcεRI^+^ hematopoietic progenitors also formed mast cells. Similar to healthy controls, population A cells from patients gave rise to mast cells following long-term culture (Fig 3G-H).

Taken together, population A cells and FcεRI^−^ hematopoietic progenitors exhibit ability to form functional mast cells. Strikingly, the cell culture assays showed that population A progenitors are more primed towards the mast cell lineage compared with FcεRI^−^ progenitors, in agreement with the gene expression profile of the primary cells (Fig 2).

### IL-3 and IL-5 induce disparate proliferative and survival promoting effects on mast cell progenitors

The culture assays showed that population A cells constituted mast cell progenitors. Whether cytokines can stimulate or modulate the phenotype of primary mast cell progenitors is largely unexplored. To identify cytokines that exhibit potential to influence mast cell progenitor functions, we plotted the gene expression patterns of selected cytokine receptors that are known to maintain hematopoietic progenitors or promote their differentiation (Fig 4). Mast cell progenitors expressed the IL-3 receptor α (*IL3RA*) and IL-5 receptor α (*IL5RA*) chains, but no or low expression of the GM-CSF receptor α (*CSF2RA*) chain. The cytokine receptor common subunit β, *CSF2RB*, which forms heterodimers with the above-mentioned α chains, was also expressed. Expression of the erythropoietin receptor (*EPOR*) was present in mast cell progenitors whereas thrombopoietin receptor (*MPL*), and FLT-3 (*FLT3*) was low or undetectable. Notably, we also observed specific expression of the IL-33 receptor, *IL1RL1*, in mast cell progenitors.

**Fig 4.**
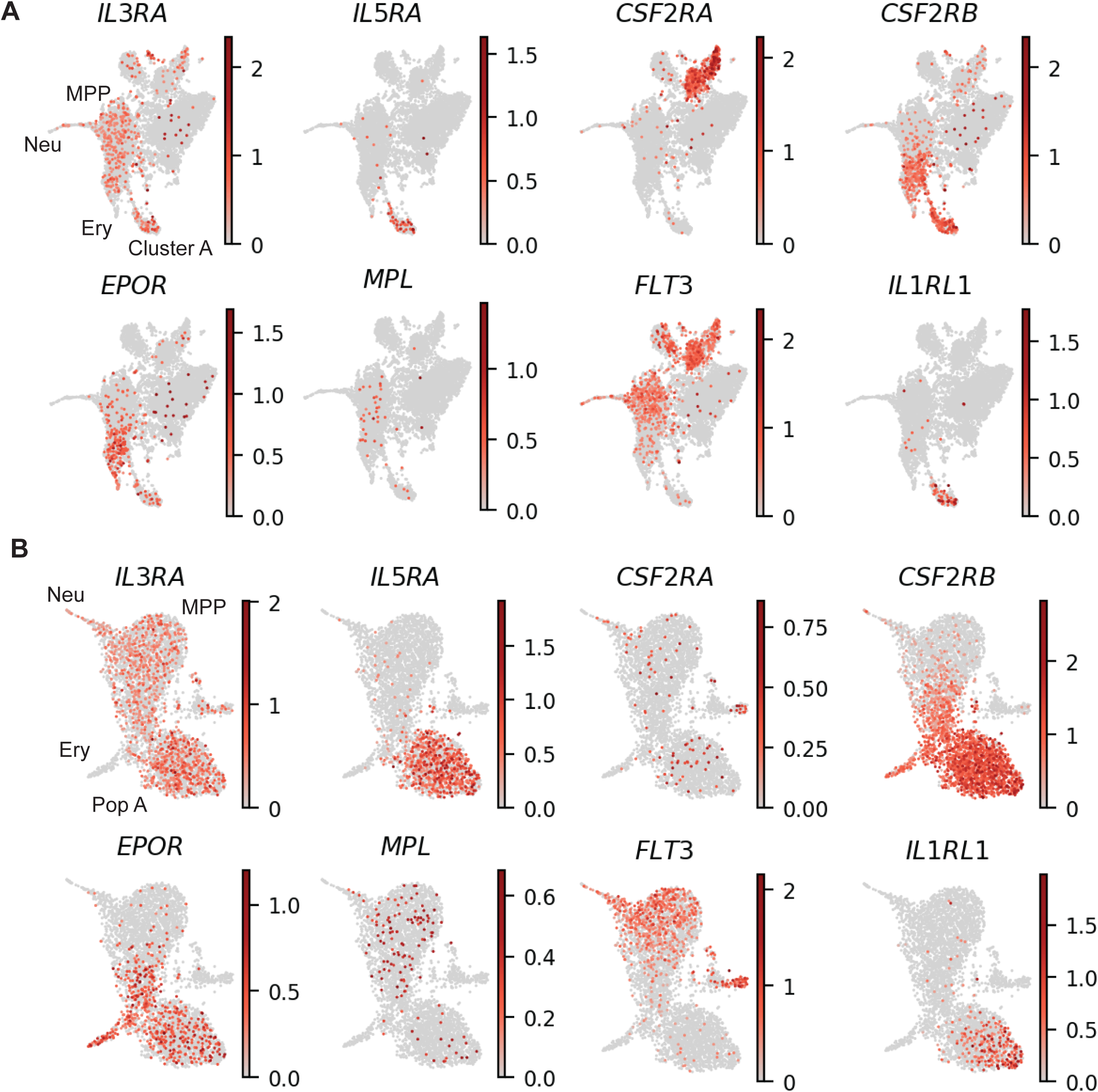
Mast cell progenitors express transcripts of the receptors for IL-3, IL-5, EPO and IL-33. The graphs show gene expression levels of the indicated cytokine receptors. A) UMAP visualization of Lin^−^ c-Kit^+^ cells (dataset 1). B) UMAP visualization of Lin^−^ CD34^+^ c-Kit^+^ cells (dataset 2).

Next, we explored the effects of the corresponding ligands – IL-3, IL-5, GM-CSF, EPO, TPO, Flt3L and IL-33 – on primary mast cell progenitors. Hematopoietic progenitor cells were isolated by magnetic-activated cell sorting and then stimulated with the individual cytokines. Flow cytometry analysis quantified the frequency of c-Kit^hi^ FcεRI^+^ cells (Fig 5A). In some experiments, the hematopoietic progenitors were labeled with CellTrace Far Red to enable analysis of the proliferative response. Treatment with IL-3 resulted in robust maintenance of c-Kit^hi^ FcεRI^+^ cells, and this condition was therefore used as reference in all experiments. Of the other cytokines analyzed, only IL-5 maintained the c-Kit^hi^ FcεRI^+^ cell population above background levels (Fig 5B).

**Fig 5.**
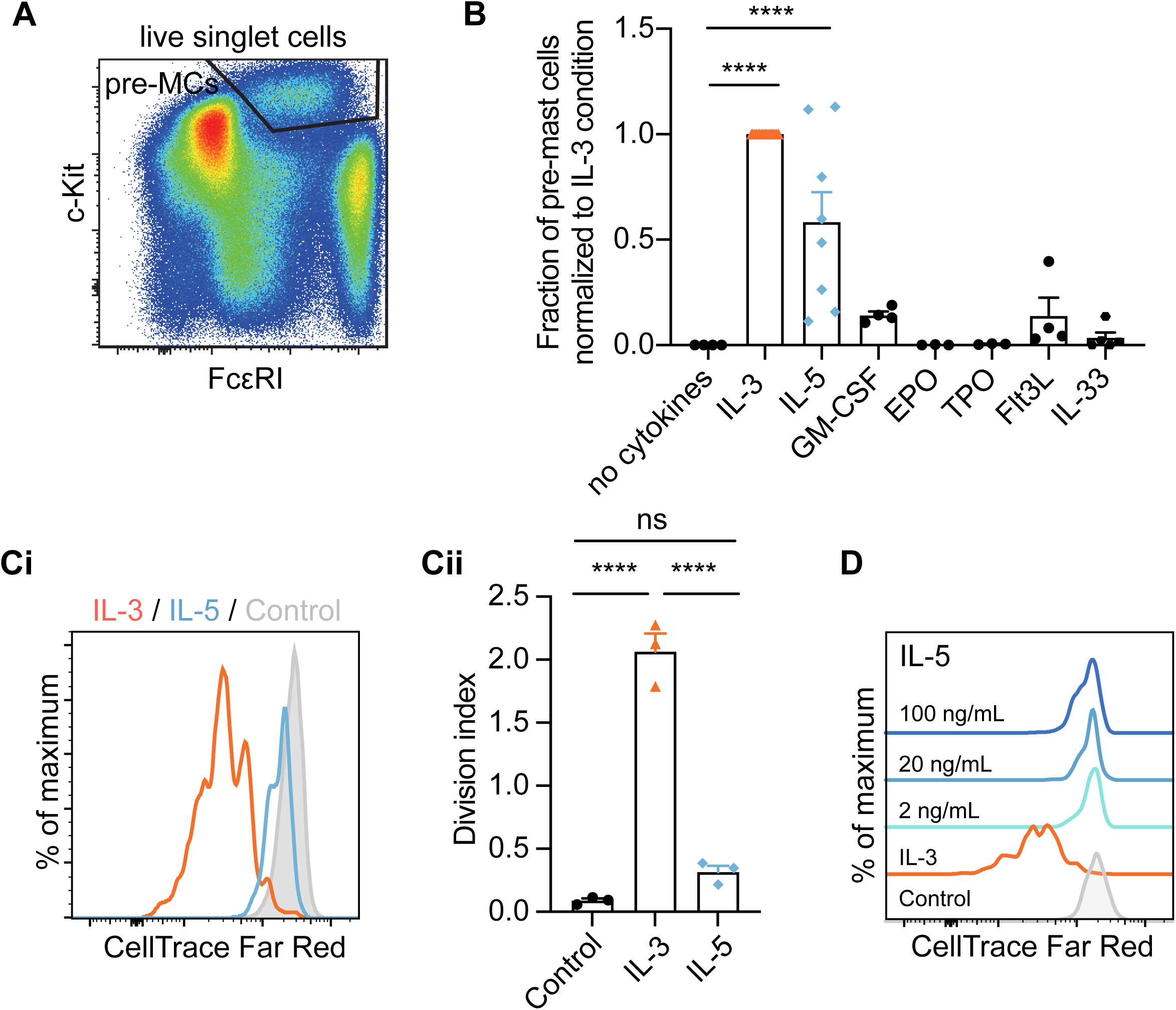
IL-3 promotes mast cell progenitor proliferation whereas IL-5 supports mast cell progenitor survival. A) Gating of pre-mast cells (pre-MCs) following culture of c-Kit^+^ progenitors for 5 days. B) Quantification of pre-mast cells following culture of c-Kit^+^ progenitors in various cytokine conditions. Points in each group represent individual buffy coats. Values are normalized to the IL-3 condition of each donor. One-way ANOVA, Dunnett’s posthoc with no cytokines as control group. C) The proliferative response of mast cell progenitors was analyzed by CellTrace Far Red signal in pre-mast cells (n=3). The control refers to analysis of all live cells following culture without cytokines, as pre-mast cells were virtually absent. D) Analysis of the proliferative response of mast cell progenitors to various concentrations of IL-5. IL-3 served as positive control. Representative of 2 independent experiments. 10 ng/ml IL-3, 20 ng/ml IL-5, 10 ng/ml GM-CSF, 1 U/ml erythropoietin (EPO), 50 ng/ml thrombopoietin (TPO), 50 ng/ml Flt3L, 22.5-100 ng/ml IL-33 unless otherwise specified. \*\*\*\**P*<0.0001.

CellTrace Far Red staining revealed that IL-3 induced strong proliferation of mast cell progenitors (Fig 5C). By contrast, IL-5 failed to stimulate robust cell proliferation (Fig 5C). We titrated the IL-5 concentration to investigate whether increasing concentrations of IL-5 promoted mast cell progenitor proliferation. However, concentrations up to 100 ng/ml IL-5 failed to induce a strong proliferative response (Fig 5D). The proliferation-promoting effect of IL-3 was concentration-dependent with low concentrations inducing little proliferation, as expected (Fig S4).

Taken together, the results show that IL-3 and IL-5 exhibit disparate proliferative and survival-promoting effects on the mast cell progenitors.

### IL-33 downregulates FcεRI expression on mast cell progenitors

Most cytokines investigated failed to maintain a high fraction of c-Kit^hi^ FcεRI^+^ cells following in vitro culture (Fig 5B). However, the cytokines investigated can potentially regulate pathways unrelated to survival and proliferation, which include alteration of the cell surface phenotype. Given the debate on whether mast cell progenitors express FcεRI, we specifically analyzed whether any cytokine shows potential to downregulate FcεRI on primary mast cell progenitors. We cultured hematopoietic progenitors with IL-3 plus one additional cytokine to maintain precursors of mast cells in culture, while simultaneously enabling analysis of individual cytokines’ effects on the FcεRI expression. IL-5, GM-CSF, EPO, TPO and Flt3L failed to modulate the FcεRI expression (Fig 6A). However, IL-33 induced strong downregulation of FcεRI on mast cell progenitors (Fig 6A). To explore whether IL-33 exerted a direct effect on mast cell progenitors, we first FACS isolated mast cell progenitors and subjected them to IL-3 alone or in combination with IL-33 (Fig 6B-C). Addition of IL-33 caused robust FcεRI downregulation, consistent with the results in which c-Kit^+^ cells were stimulated in bulk (Fig 6A,C). Taken together, exposure to IL-33 leads to FcεRI downregulation in primary mast cell progenitors.

**Fig 6.**
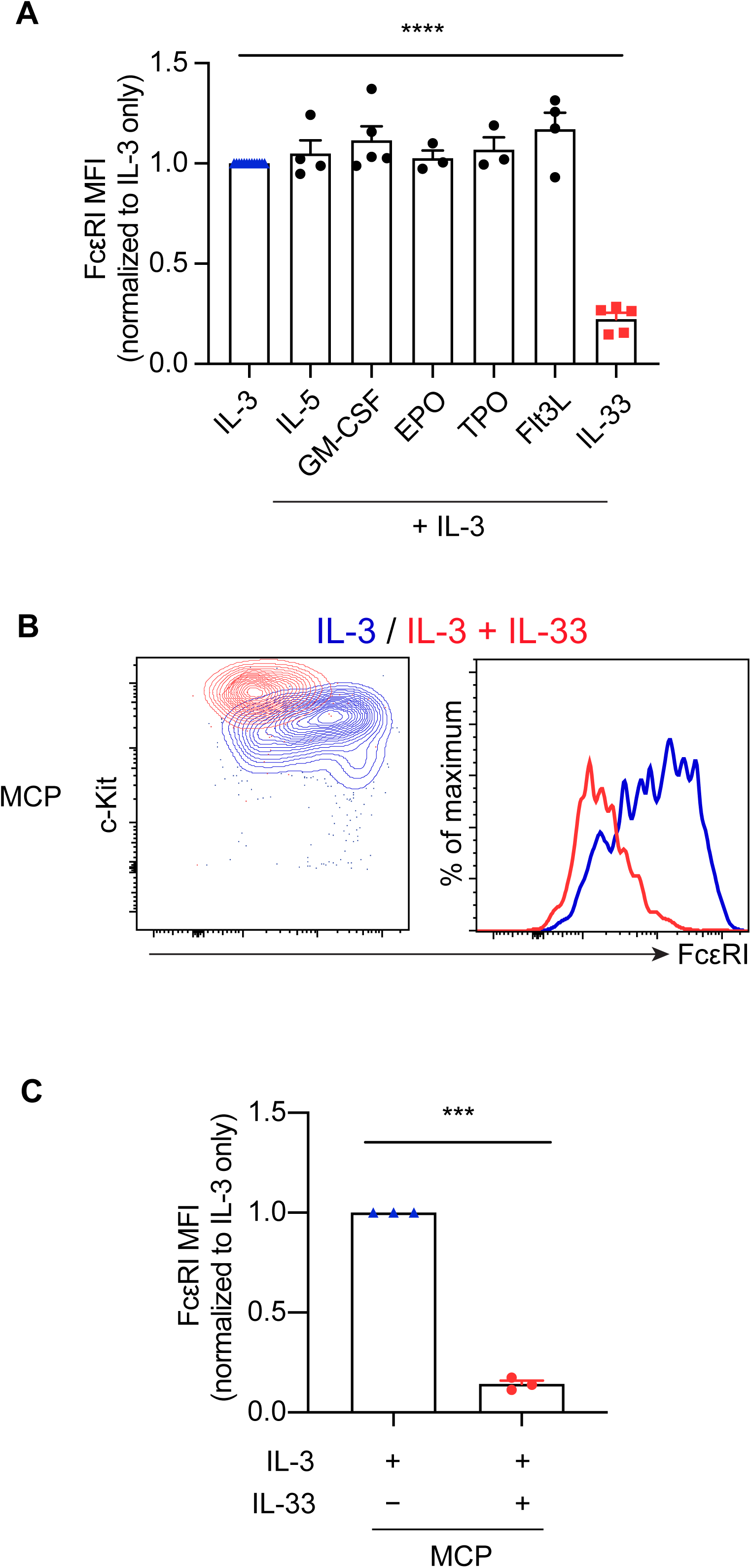
IL-33 downregulates Fc Ε RI expression on mast cell progenitors. A)Quantification of Fc Ε RI expression on pre-mast cells following 5 days in culture with the indicated cytokines. Points in each group represent individual buffy coats. The levels were normalized to the IL-3 condition of each donor. 10 ng/ml IL-3, 20 ng/ml IL-5, 10 ng/ml GM-CSF, 1 U/ml erythropoietin (EPO), 50 ng/ml thrombopoietin (TPO), 50 ng/ml Flt3L, 22.5-100 ng/ml IL-33 unless otherwise specified. B) Flow cytometry analysis of FACS-isolated mast cell progenitors (MCPs) cultured with 10 ng/ml IL-3 alone or in combination with 100 ng/ml IL-33 for 5 days. Cultures of Fc Ε RI^−^ progenitors are shown in Fig S5. C) Quantification of the panel B (n=3). MFI, median fluorescent intensity. Two-tailed one-sample t-test in panels A and C, hypothetical value 1.0. \*\*\**P*<0.001, \*\*\*\**P*<0.0001.

## Discussion

The single-cell transcriptional landscapes presented here suggest that Lin^−^ CD34^+^ c-Kit^+^ cells in blood constitute a spectrum of hematopoietic progenitors and that there is a specific branch of FcεRI^+^ progenitors. It is expected that FcεRI^−^ hematopoietic progenitors exhibit mast cell-forming capacity (*5*), given the presence of multipotent progenitor-like cells in the landscapes. However, the observation that FcεRI^−^ hematopoietic progenitors give rise to mast cells does not rule out the existence of FcεRI^+^ mast cell progenitors. In agreement with this, we show that FcεRI^+^ progenitors give rise to mast cells in long-term culture assays. This challenges the idea that FcεRI upregulation is a late event during mast cell maturation.

FcεRI expression has been assumed to increase with mast cell maturation (*3*). Aberrant mast cells without FcεRI expression are observed in a subset of patients with systemic mastocytosis, contributing to the general view that mast cells originate from FcεRI^−^ progenitors. The inability to detect CD34^+^ FcεRI^+^ progenitors in a study of systemic mastocytosis patients has strengthened this view further (*5*). We confirm that FcεRI^−^ progenitors in patients exhibit mast cell-forming capacity. However, we were able to isolate a population of FcεRI^+^ progenitors from systemic mastocytosis patients, and these cells gave rise to mast cells. Thus, FcεRI^+^ progenitors constitute a potential source of mast cells also in patients with systemic mastocytosis. The inability to detect FcεRI^+^ progenitors in a previous report of systemic mastocytosis patients and normal controls is likely attributed to the difference in the method to enrich for hematopoietic progenitors (*5*). We used c-Kit-based cell selection, as the CD34 levels on mast cell progenitors is lower than that of the main hematopoietic progenitor population. An affinity-based CD34 isolation strategy is likely to enrich for cells with the highest CD34 levels, which can lead to failure in detecting the mast cell progenitor population. Another possible explanation to the lack of FcεRI on mast cell progenitors is if the microenvironment from which the cells are extracted promotes FcεRI downregulation. However, both studies used peripheral blood as a source and included healthy controls and systemic mastocytosis patients. IL-33-mediated downregulation of FcεRI or subject-related differences are therefore unlikely to explain the discrepancy. This favors the idea that technical differences can explain the inability to detect FcεRI^+^ progenitors.

We show that the bulk of the FcεRI^−^ progenitors in peripheral blood at steady-state conditions are less primed to the mast cell lineage than FcεRI^+^ progenitors. However, the observation that FcεRI levels discriminate stages of mast cell priming in primary hematopoietic progenitors does not necessarily mean that FcεRI levels constitute a general marker of mast cell differentiation. In fact, FcεRI^−^ mast cells are observed at specific anatomical compartments (*6*). Mast cells lacking FcεRI expression are also produced *in vitro* upon differentiation of fetal liver and cord blood progenitor cells, and the commonly used mast cell line HMC-1 lack FcεRI expression (*21-24*). Experiments showing that external factors can regulate FcεRI levels in mast cells past the progenitor stage further supports the idea that FcεRI expression is not immediately linked with mast cell differentiation (*22, 25, 26*).

Studies have shown that CD34 downregulation may precede FcεRI upregulation during mast cell differentiation in vitro (*4, 5*). Here, we demonstrate that CD34 and FcεRI expression may co-exist on single mast cell progenitors. These observations are not necessarily contradictory. Instead, they suggest that we need to revise the model in which FcεRI expression alone is a marker of mast cell differentiation. However, differentiation along the mast cell differentiation trajectory in hematopoietic progenitors is associated with increased FcεRI expression in steady-state hematopoiesis.

The culture assay presented here was designed to identify cytokines with major and specific effects on mast cell progenitor survival or phenotype. We cannot exclude the possibility that cytokines that failed to maintain a pre-mast cell population indeed promote mast cell progenitor survival or exhibit regulatory functions. For example, cytokines known to promote survival, proliferation and differentiation of non-mast cell lineage cells might obscure survival-promoting effects on mast cell progenitors. It is also possible that the cytokines regulate pathways that were not analyzed. Nevertheless, we revealed that IL-3, IL-5 and IL-33 exhibit proliferative, survival-promoting and FcεRI regulatory functions, respectively, on primary mast cell progenitors.

Survival-promoting functions of cytokines have been investigated on in vitro-derived mature mast cells (*27, 28*). Yet, the regulatory effects of cytokines on primary mast cell progenitors are largely unexplored. Ochi et al performed thymidine incorporation assays to study the proliferative effects of individual cytokines on cord blood-derived immature mast cells (*28*). In line with our results, Ochi et al revealed proliferation-promoting effects of IL-3 on in vitro-derived immature mast cells and IL-5 did not induce proliferation of the immature mast cells (*28*). We showed that while IL-5 is insufficient to promote significant mast cell progenitor proliferation, it is able to promote cell survival.

IL-3 and IL-5 share a common β chain for signaling (*29*). Nevertheless, these cytokines promoted disparate effects on the mast cell progenitors at the initial concentrations tested. Increasing doses of IL-5 failed to induce a proliferative response similar to the capacity of IL-3. A limited number of IL-5 receptors on the cell surface could provide a possible explanation. However, it has yet to be determined whether the α chains of the receptors are involved in cell signaling, which could instruct disparate signaling pathways (*29*).

The single-cell transcriptomics datasets presented with this investigation provide the research community with a high-resolution map of hematopoietic progenitor differentiation with focus on the rare mast cell progenitor population. We show how the data can be used to reveal cell lineage bias and factors that regulate mast cell progenitors. Notably, we reveal a temporal association between mast cell lineage differentiation and the upregulation of FcεRI in hematopoietic progenitors. These data provide definitive proof that FcεRI may appear already at the mast cell progenitor stage, before terminal mast cell maturation.

## Methods

### Study design

The study aimed to chart hematopoiesis with focus on mast cell differentiation, specifically the regulation of FcεRI expression. The exploratory design of the study, including the generation of single-cell RNA sequencing data of hematopoietic progenitors, was complemented with validation experiments and functional assays.

### Ethics statement

The Swedish Ethical Review Authority and the Regional Review Board in Stockholm approved the study and the subjects provided informed consent to participate in the study.

### Cell isolation, culture and analysis

Cells from healthy donors were extracted from donated buffy coat fractions of peripheral blood, whereas cells from systemic mastocytosis patients were extracted from peripheral blood. Results in the manuscript are derived from analysis of healthy donors unless otherwise stated. Ficoll separation isolated blood mononuclear cells from the buffy coats. c-Kit^+^ cells were extracted with the CD117 Microbead Kit (Miltenyi Biotec, Bergisch Gladbach, Germany). Cells were cultured in complete StemPro-34 serum-free medium (Gibco/Thermo Fisher Scientific, Waltham, MA) with 2 mM L-glutamine (Cytiva Hyclone, Global Life Sciences Solutions USA, Marlborough, MA), 100 U/ml penicillin and 0.1 mg/ml streptomycin (Cytiva Hyclone). The recombinant cytokines IL-3, IL-6, Thrombopoietin, IL-33, GM-CSF, Flt3L (all from Peprotech, Rocky Hill, NJ), IL-5 (Peprotech and R&D Systems, Minneapolis, MN), erythropoietin (Janssen-Cilag Ab, Solna, Sweden), and stem cell factor (Swedish Orphan Biovitrum, Stockholm, Sweden) supplemented the medium. Cytokines concentrations are specified in the figures and figure legends. In selected experiments, cells were starved from stem cell factor overnight before flow cytometry analysis to prevent c-Kit internalization and promote c-Kit upregulation (*30*). Cytocentrifugation was performed using Shandon Octospot (Thermo Fisher Scientific) where the cells numbers were limited. Tryptase staining was performed as described using Z-Gly-Pro-Arg-4-Methoxy-β-naphthylamide substrate (*31*). May-Grünwald and Giemsa solutions (Sigma-Aldrich/Merck, Munich, Germany) stained the cytocentrifuged cells.

### Flow cytometry

Antibodies used for flow cytometry analysis included CD3 (SK7), CD14 (M5E2), CD19 (HIB19), CD15 (W6D3), CD34 (581), CD63 (H5C6), CD117 (A3C6E2 and 104D2), FcεRI (CRA-1) and β2-microglobulin (2M2). TotalSeq-B0953 PE Streptavidin was used to visualize β2-microglobulin-biotin labeling during FACS and single-cell transcriptomics analysis. The viability dyes DAPI and 7-AAD were used where appropriate (BD and Thermo Fisher Scientific). CellTrace Far Red (Thermo Fisher Scientific) served as marker of proliferation. The Canto II or LSRFortessa flow cytometer was used to analyze cells (BD Biosciences, San Jose, CA). Fluorescence-activated cell sorting on the FACS Aria Fusion system (BD Biosciences) isolated specific cell subsets. The FlowJo software (TreeStar, Ashland, OR) was used to analyze the data.

### Single-cell transcriptomics analysis

Lin^−^ c-Kit^+^ cells were isolated with FACS and processed using the Chromium Single Cell 3’ Reagents Kit v2 (10X Genomics, Pleasanton, CA). The library was sequenced on the NovaSeq 6000 SP flow cell with the 26-8-0-96 read setup. An antibody hashtag-based approach distinguished the single-cell transcriptomes of Lin^−^ CD34^+^ c-Kit^+^ and Lin^−^ CD34^+^ c-Kit^+^ FcεRI^+^ hematopoietic progenitors. Briefly, a sample with MACS-enriched c-Kit^+^ cells was divided into two fractions. One fraction was stained with biotinylated β2-microglobulin antibody and the other fraction with unlabeled β2-microglobulin antibody. Each cell fraction was then incubated with flow cytometry antibodies plus oligonucleotide-labeled streptavidin-PE. Lin^−^ CD34^+^ c-Kit^+^ cells without the hashtag (PE^−^) and Lin^−^ CD34^+^ c-Kit^+^ FcΕRI^+^ cells with the hashtag (PE^+^) were FACS sorted. The sorted cells were pooled and processed using the Chromium Next GEM Single Cell 3’ Reagent Kit v3.1 (dual index) with Feature Barcode technology for Cell Surface Protein (10X Genomics). The gene expression and antibody libraries were sequenced on the NovaSeq 6000 SP flow cell with the 28-10-10-90 read setup. The Cell Ranger (Version 3.0.1 and 5.0.1 for dataset 1 and 2, respectively) pipeline and Scanpy (*32*) were used to process the sequencing data.

The Cell Ranger pipeline identified 7336 Lin^−^ c-Kit^+^ cells (dataset 1). The following preprocessing steps were applied: filtering out cells expressing less than 500 genes, cells with more than 5% mitochondrial gene content, cells with more than 4000 total genes with at least one count, and genes expressed in fewer than 3 cells. The pre-processing recipe from Zheng et al (2017) (*33*) was applied and the data was fed to Scrublet (*34*) for doublet simulation and removal of 200 doublet cells. The raw expression of the remaining 6874 cells was then re-processed with the above recipe, with the highly variable genes screened through a correlation filter for cell cycle genes (cell cycle gene list downloaded from GSEA, MSigDB version 7.0, August 2019, entry: REACTOME_CELL_CYCLE, identifier: R-HSA-1640170). PCA using 50 components, a size of local neighborhood defined using 15 neighbors, and Leiden clustering (*35*) with a resolution parameter of 0.5 was used for the UMAP embedding. The gene expression layer for visualizations on the UMAP embedding was normalized per cell, followed by log1p transformation.

The Cell Ranger pipeline identified 4391 Lin^−^ CD34^+^ c-Kit^+^ cells (dataset 2). Features containing oligonucleotide-labeled streptavidin-PE counts were split from the gene expression matrix for pre-processing: filtering out cells expressing less than 500 genes, cells with more than 6% mitochondrial gene content, cells with more than 6000 total genes with at least one count, and genes expressed in fewer than 3 cells. The pre-processing recipe from Zheng et al (2017) (*33*) was applied and the data was fed to Scrublet (*34*) for doublet simulation, removing 99 doublets. The raw gene expression of the remaining 4063 cells was reprocessed using the above recipe, and highly correlated cell cycle genes were removed. PCA using 50 components, a size of local neighborhood defined using 15 neighbors, and Leiden clustering with a resolution parameter of 0.9 was used for the UMAP embedding, for a total of 12 clusters. The gene expression layer for visualizations on the UMAP embedding was normalized per cell, followed by log1p transformation. The oligonucleotide-labeled streptavidin-PE antibody features were adjusted for log1p values and mapped to the UMAP. Diffusion pseudotime (*36*) was calculated using the MPP cluster as the root. Temporal ordering of the cells from the MPP cluster through the intermediate cluster, ending with the population A (Pop A) clusters was refined by introducing a ranking system combining the Leiden clusters with pseudotime. Cells were first ordered by cluster, then sorted by pseudotime within the cluster ordering. Identification of dynamically regulated genes was based on Tusi et al (2018) (*15*). This was conducted on all genes that passed pre-processing prior to the recipe from Zheng et al (2017) (*33*), using a rolling window of size 60, with dynamic genes defined as having adjusted *P*-values < 0.0005. A Spearman correlation was performed between the ranked order and the expression of dynamically regulated genes to identify Pop A signature genes. A correlation score cut-off of > 0.5 was applied. A rolling average of 20 was applied to visualize gene expression levels in Fig 2H and Fig 2J.

The data reported in this article have been deposited in the Gene Expression Omnibus database (GSE184351). Requests to access raw sequencing data can be submitted to the corresponding author. Permission to access raw data is provided if the conditions in the ethical permit can be met.

### Statistical analysis

Statistical analysis was performed using Prism software (Graphpad software, La Jolla, CA). *P*<0.05 was considered significant. The statistical tests used are specified in the figure legends.

## Acknowledgements

We acknowledge support from the National Genomics Infrastructure in Stockholm, funded by the Science for Life Laboratory, the Knut and Alice Wallenberg Foundation, and the Swedish Research Council. We are also grateful for support from SNIC/Uppsala Multidisciplinary Center for Advanced Computational Science for assistance with massively parallel sequencing analysis, and access to the UPPMAX computational infrastructure. We thank Bertie Göttgens at Wellcome - MRC Cambridge Stem Cell Institute, University of Cambridge for facilitating the bioinformatics analysis. Recombinant stem cell factor was a generous gift from Swedish Orphan Biovitrum.

## Authors contributions

C.W. and O.B. performed experiments. M.E., J.G. and C.S. helped with execution of experiments. C.W. analyzed the cell culture data. D.B. analyzed the single-cell transcriptomics data. J.S.U. diagnosed the patients and organized the collection of the samples. M.S.V. reviewed and contributed to the bioinformatics analysis. N.K.W. helped coordinating the bioinformatics analysis. J.S.D. and C.W. designed the experiments. G.N. and J.S.D. supervised the study. J.S.D. secured funding. J.S.D., C.W, and D.B. drafted the manuscript. All authors contributed to the final version of the manuscript.

