## Supplementary material for "Single-cell transcriptomics reveals the identity and regulators of human mast cell progenitors": Table S1

**Table S1. Population A signature genes include well-defined mast cell markers**

| Well-defined mast cell markers* | Population A signature gene |
|---|---|
| *KIT* | Yes |
| *IL1RL1* | No |
| *FCER1A* | Yes |
| *MS4A2* | Yes |
| *ENPP3* | No |
| *HDC* | Yes |
| *TPSAB1* | Yes |
| *TPSB2* | Yes |
| *TPSD1* | No |
| *CMA1* | No |
| *CPA3* | Yes |
| *CTSG* | No |
| *HPGDS* | Yes |
| *LTC4S* | Yes |

*Markers defined in E. Motakis, S. Guhl, Y. Ishizu, M. Itoh, H. Kawaji, M. de Hoon, T. Lassmann, P. Carninci, Y. Hayashizaki, T. Zuberbier, A. R. Forrest, M. Babina, F. consortium, Redefinition of the human mast cell transcriptome by deep-CAGE sequencing. *Blood* **123**, e58-67 (2014).
