## Supplementary Figures for "Single-cell transcriptomics reveals the identity and regulators of human mast cell progenitors"

### Supplementary Figure legends

**Figure S1. Plotting the expression of cell lineage-associated genes allows annotation of the Lin$^{-}$ c-Kit$^{+}$ peripheral blood cell dataset.**

The expression patterns of cell lineage-associated genes annotate dataset 1 (related to Fig 1).

**Figure S2. Gating strategies to FACS isolate cells for the generation of dataset 2.**

The graphs show the isolation strategies for Lin$^{-}$ CD34$^{+}$ c-Kit$^{+}$ cells and PE-oligonucleotide-labeled population A cells (red).

**Figure S3. Plotting the expression of cell lineage-associated genes allows annotation of the Lin$^{-}$ CD34$^{+}$ c-Kit$^{+}$ peripheral blood cell dataset.**

The expression patterns of cell lineage-associated genes annotate dataset 2 (related to Fig 2).

**Figure S4. Decreasing levels of IL-3 result in diminished mast cell progenitor proliferation.**

MACS-isolated c-Kit$^{+}$ progenitors cultured for 5 days were analyzed with flow cytometry and pre-mast cells were gated. A) Proliferative response measured with CellTrace Far Red to decreasing concentrations of IL-3. The control refers to analysis of all live cells following culture without cytokines, as pre-mast cells were virtually absent. B) Quantification of the results in panel B (n=3-4 donors per concentration). One-way ANOVA with Dunnett's multiple comparisons test relative to the control group. ****$P$<0.0001.

**Figure S5. Analysis of FcεRI$^{-}$ progenitors cultures provides a reference for mast cell progenitor cultures.**

Flow cytometry analysis of FACS-isolated FcεRI$^{-}$ progenitors cultured with 10 ng/ml IL-3 alone or in combination with 100 ng/ml IL-33 for 5 days. Related to Fig 6B.

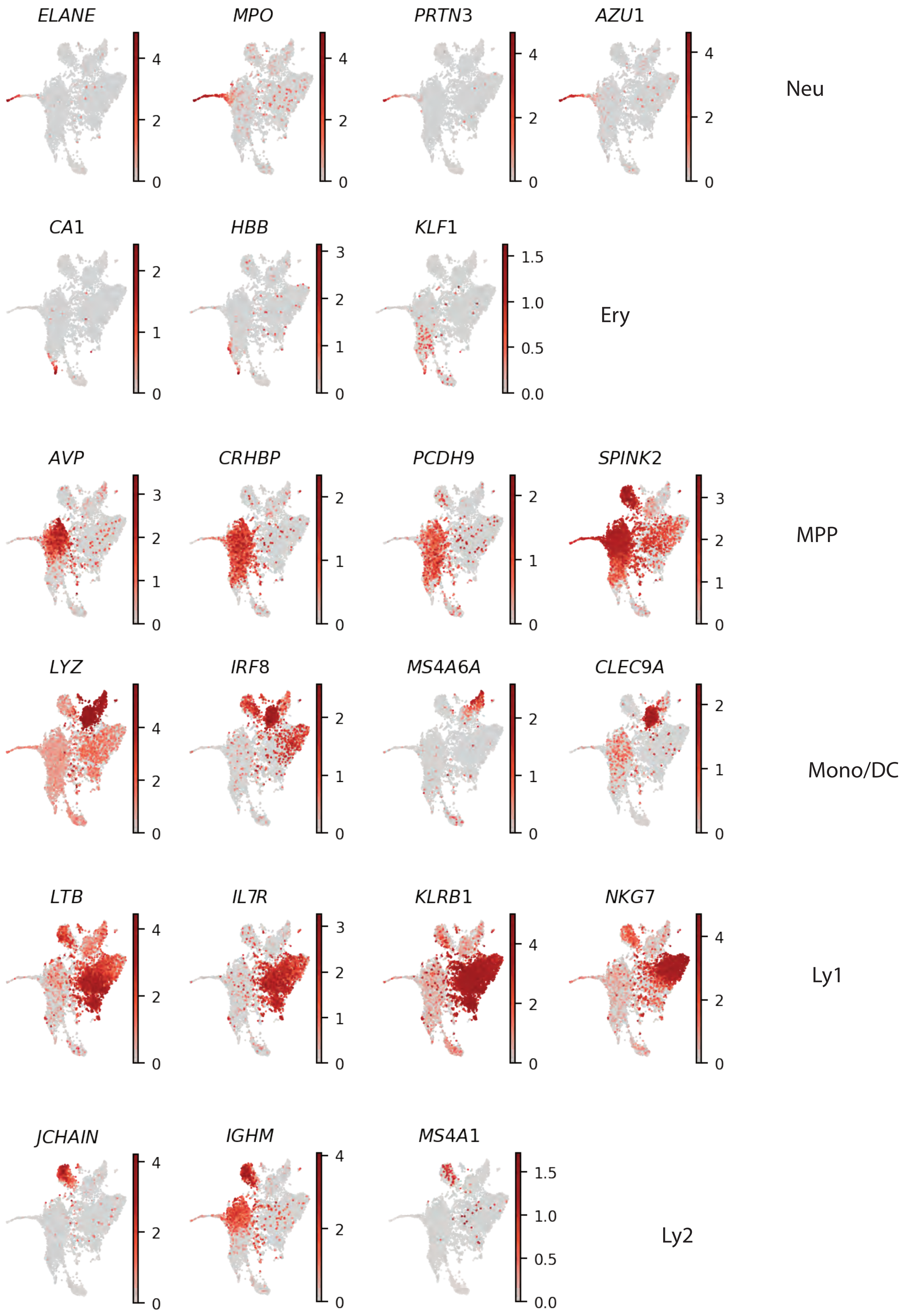

ELANE
MPO
PRTN3
AZU1
Neu
CA1
HBB
KLF1
Ery
AVP
CRHBP
PCDH9
SPINK2
MPP
LYZ
IRF8
MS4A6A
CLEC9A
Mono/DC
LTB
IL7R
KLRB1
NKG7
Ly1
JCHAIN
IGHM
MS4A1
Ly2


**Figure S1**

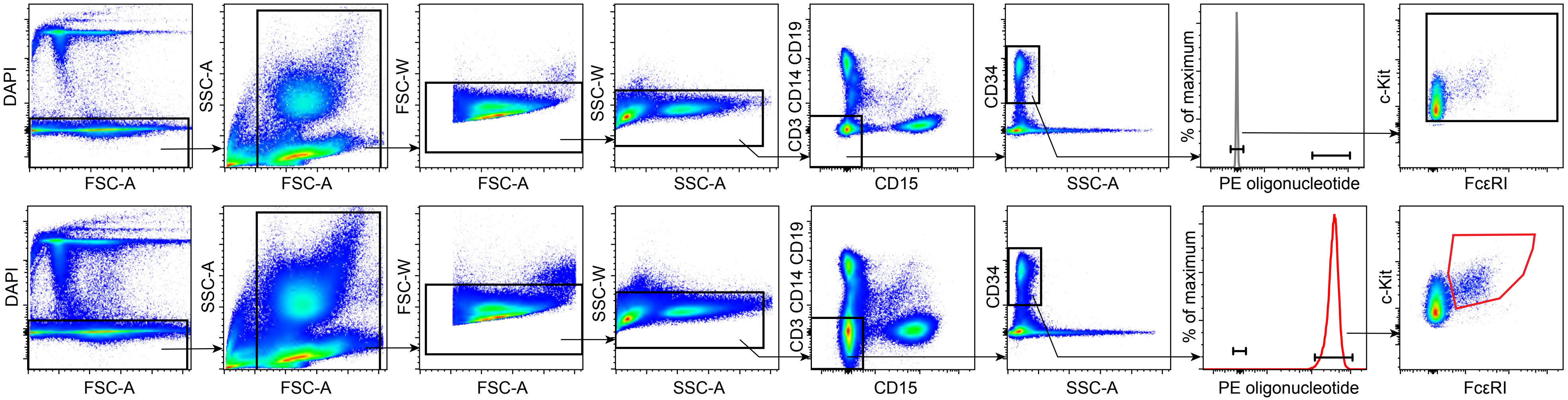
DAPI
FSC-A
SSC-A
FSC-A
FSC-W
FSC-A
SSC-W
SSC-A
CD3 CD14 CD19
CD15
CD34
SSC-A
% of maximum
PE oligonucleotide
c-Kit
FcεRI
DAPI
FSC-A
SSC-A
FSC-A
FSC-W
FSC-A
SSC-W
SSC-A
CD3 CD14 CD19
CD15
CD34
SSC-A
% of maximum
PE oligonucleotide
c-Kit
FcεRI

**Figure S2**

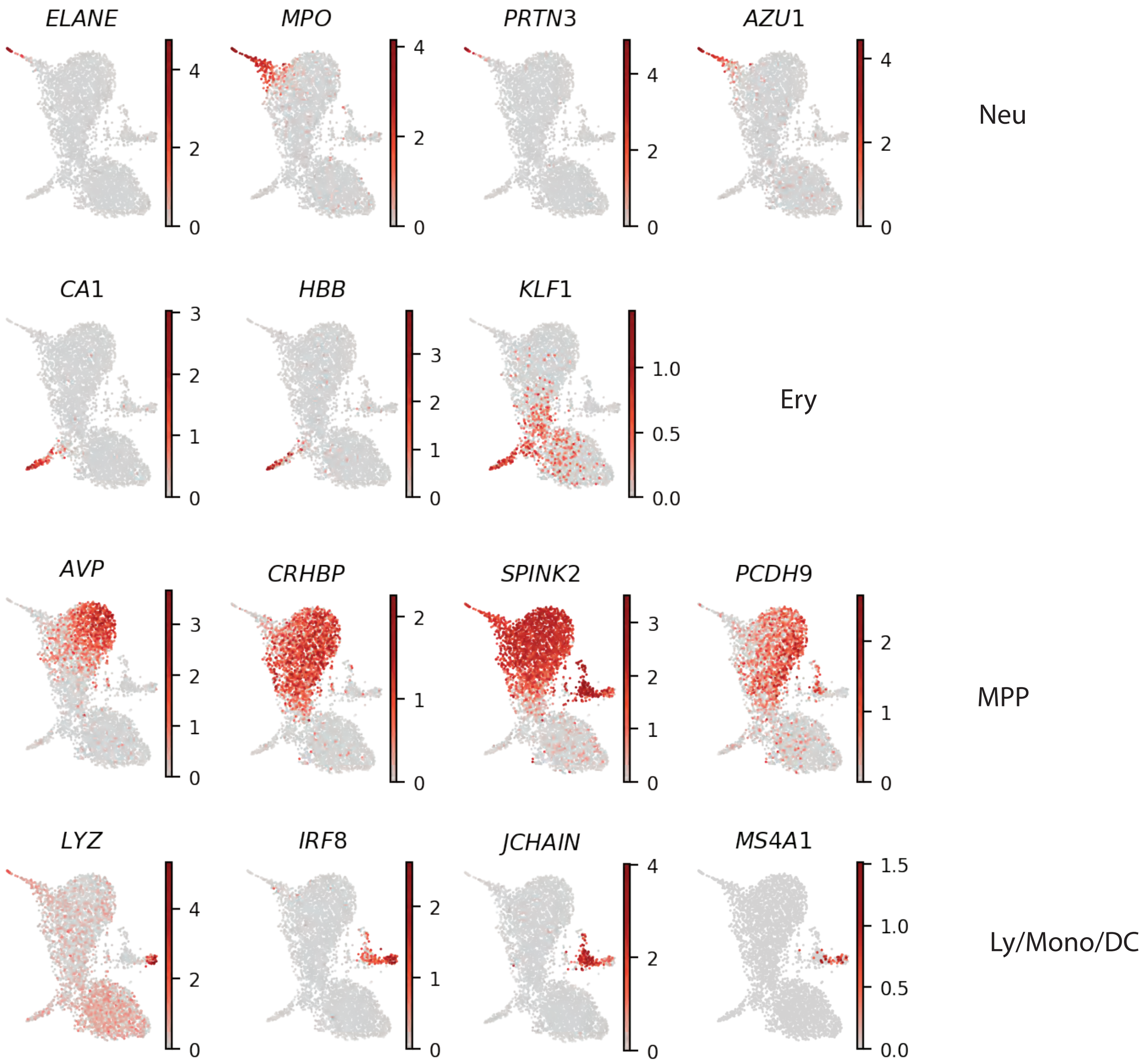

ELANE
MPO
PRTN3
AZU1
Neu
CA1
HBB
KLF1
Ery
AVP
CRHBP
SPINK2
PCDH9
MPP
LYZ
IRF8
JCHAIN
MS4A1
Ly/Mono/DC


**Figure S3**

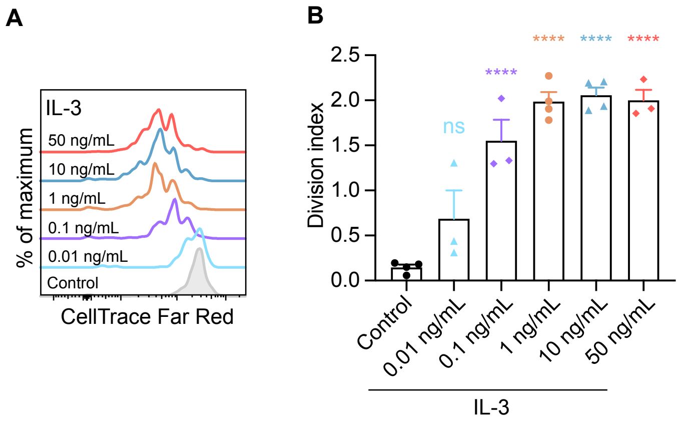

A
IL-3
50 ng/mL
10 ng/mL
1 ng/mL
0.1 ng/mL
0.01 ng/mL
Control
% of maximum
CellTrace Far Red
B
Division index
2.5
2.0
1.5
1.0
0.5
0.0
ns
****
****
****
****
Control
0.01 ng/mL
0.1 ng/mL
1 ng/mL
10 ng/mL
50 ng/mL
IL-3


**Figure S4**

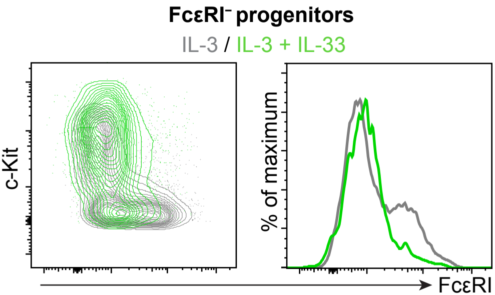


**Figure S5**
